# YME1L-dependent regulation of mitochondrial Ca^2+^ transport: a role for mitochondrial uptake protein 1 (MICU1) in nutrient sensing

**DOI:** 10.64898/2025.12.11.693667

**Authors:** Julian D. C. Serna, Donato D’Angelo, Georgia Ohya, Rosario Rizzuto, Alicia J. Kowaltowski, Anna Raffaello

**Affiliations:** Department of Biomedical Sciences, University of Padova, Padova, Italy; Departamento de Bioquímica, Instituto de Química, Universidade de São Paulo, São Paulo, Brazil; National Center on Gene Therapy and RNA-Based Drugs, Padova, Italy; Myology Center (CIR-Myo), University of Padova, Padova, Italy

## Abstract

Mitochondrial calcium uptake via the mitochondrial calcium uniporter complex (MCUc) is tightly regulated by gatekeeper proteins such as MICU1 and MICU2. While long-term nutritional interventions have been shown to remodel the MCUc, its short-term regulation during postprandial transitions and acute nutrient stress remains unclear. Here, we demonstrate that non-gated MCUc increases in the postprandial liver due to a transient loss of mature MICU1 (m-MICU1), accompanied by changes in its precursor form (p-MICU1). These changes are more pronounced and occur earlier than the general increase in mitochondrial mass and respiratory protein content, indicating that they are independent of mitochondrial remodelling during refeeding.

*In vitro*, serum deprivation, insulin signalling, modulation of mTOR activity, and glutamine starvation all affect mature or precursor MICU1 protein levels, indicating that multiple nutritional changes promote MICU1 processing alterations, leading to the formation of a non-gated, MICU-deficient MCUc. Under these conditions, mitochondrial, but not cytosolic, Ca²⁺ levels increase in intact cells. Moreover, mitochondrial Ca²⁺ uptake rates are enhanced, which correlates with higher mitochondrial oxygen consumption, increased reactive oxygen species (ROS) production, and enhanced sensitivity to mitochondrial permeability transition.

We identify the mitochondrial protease YME1L as essential for MICU1 turnover in response to nutrient availability. YME1L silencing prevents MICU1 degradation and suppresses the enhanced Ca²⁺ uptake induced by glutamine deprivation. Together, these results reveal nutrient-sensitive, YME1L-dependent remodelling of the MCU complex and establish MICU1 degradation as a key mechanism linking metabolic signals to mitochondrial Ca²⁺ transport.

## Introduction

Cell metabolism must adapt to nutritional variations ^1, 2, 3^. This metabolic flexibility largely depends on the ability of mitochondria to sense and respond to environmental stimuli. Changes in mitochondrial Ca^2+^ concentrations ([Ca^2+^]_mit_) enable these organelles to adjust their function according to cellular demands ^4^. Small and transient increases in [Ca^2+^]_mit_ enhance oxidative metabolism by activating substrate transporters and matrix dehydrogenases ^4, 5^. Moreover, mitochondrial Ca^2+^ plays a critical role in regulating preferences for substrate oxidation, which significantly impact both cellular and whole-organism metabolism ^6^.

Alterations in mitochondrial morphology, including network shape, cristae structure, and contact sites between the endoplasmic reticulum (ER) and mitochondria, have been recognized as mechanisms regulating mitochondrial function during nutrient sensing ^1, 7^. Recently, changes in the abundance and composition of the mitochondrial Ca^2+^ uniporter complex (MCUc) and the Na^+^/Ca^2+^ exchanger (NCLX) have emerged as additional regulatory layers controlling mitochondrial function ^3, 8, 9, 10, 11^. Furthermore, TMEM65, a mitochondrial inner membrane protein, has been identified as a crucial factor in mitochondrial Ca^2+^ efflux, acting either as a positive regulator of NCLX-mediated Na^+^/Ca^2+^ exchange ^12^ or potentially functioning as an exchanger itself ^13, 14^.

The metazoan MCUc comprises MCU ^15, 16^, the essential MCU regulator (EMRE) ^17, 18^, and mitochondrial Ca^2+^ uptake proteins MICU1 and MICU2 ^19, 20, 21^. MICU1 and MICU2, along with MICU3 (expressed in muscle and the central nervous system), regulate the kinetics and threshold of Ca^2+^ uptake in a non-redundant manner ^19, 20, 21, 22, 23^. MCUb, the dominant negative isoform of MCU, reduces the permeability of the MCUc to Ca^2+^ ^24^. The MICU1/MICU2 ratio to MCU modulates tissue-specific decoding of cytoplasmic Ca^2+^ signals by mitochondria ^25^. In skeletal muscle, an MCUc containing MICU1.1/MICU2 dimer exhibits a lower threshold for Ca^2+^ activation and is essential for mitochondrial responsiveness to the functional demands of myofibers during muscle contraction ^26^.

While structural and functional roles of MCUc components are increasingly understood, less is known about cell signalling events regulating the complex. During myogenesis, RBFOX2 controls alternative splicing of the MICU1 transcript to generate MICU1.1 ^27^. At the transcriptional level, Sp1 and EGR regulate MICU1 expression ^28^. Phosphorylation of MICU1 by Akt promotes its degradation and stimulates Ca^2+^ uptake by disrupting MICU1/MICU2 control ^29^, although the protease responsible for this degradation remains unidentified.

Interestingly, MICU2 is specifically downregulated in animals subjected to 6 months of caloric restriction ^10^. Similarly, MCUb is upregulated after 48 hours of fasting, reducing mitochondrial Ca^2+^ levels and promoting lipid oxidation in muscle ^3^. These findings indicate nutrient sensitivity of the MCUc under chronic nutritional interventions, although the mechanisms controlling the MCUc components in response to nutrient availability remain unclear. In this study, using a postprandial liver model transitioning from fasting to feeding and several *in vitro* nutrient availability models, we demonstrate that MICU1 and TMEM65 undergo acute changes. This change is mediated by YME1L, which is required for the degradation of MICU1 and TMEM65 induced by nutrient deprivation. Collectively, our results highlight the role of MICU1/MICU2 and YME1L in cellular adaptations to changes in nutritional status.

## Results

### Non-gated MCUc increases as a consequence of MICU1 downregulation in the postprandial liver

The mitochondrial calcium uniporter complex (MCUc) is known to undergo remodelling in response to nutrient scarcity, including a decrease in MICU2 in animals subjected to 6 months of caloric restriction ^10^ and an upregulation of MCUb after 48 hours of fasting ^3^. To determine whether MCUc remodelling also occurs during adaptation to short-term nutrient scarcity, we established a postprandial model (**Fig. 1A**) by monitoring fasting-feeding transitions over time in mouse livers, as previously described by Sood and coworkers ^30^. The nutritional state was assessed by measuring the phosphorylation of ribosomal protein S6 (S6) at Serine 235/236 (p-S6), a marker of mTORC1 activity ^31^. As expected, p-S6 levels increased upon feeding and declined rapidly during the postprandial phase (**Fig. 1B**, quantified in **Fig. S1A**). Consistent with previous findings ^30^, the amount of cleaved OPA1 (s-OPA1) and its C-terminal fragment 2 (CTF2) increased during the postprandial period (**Fig. 1A, S1B and S1C**), validating our fasting/feeding model.

**Figure 1.**
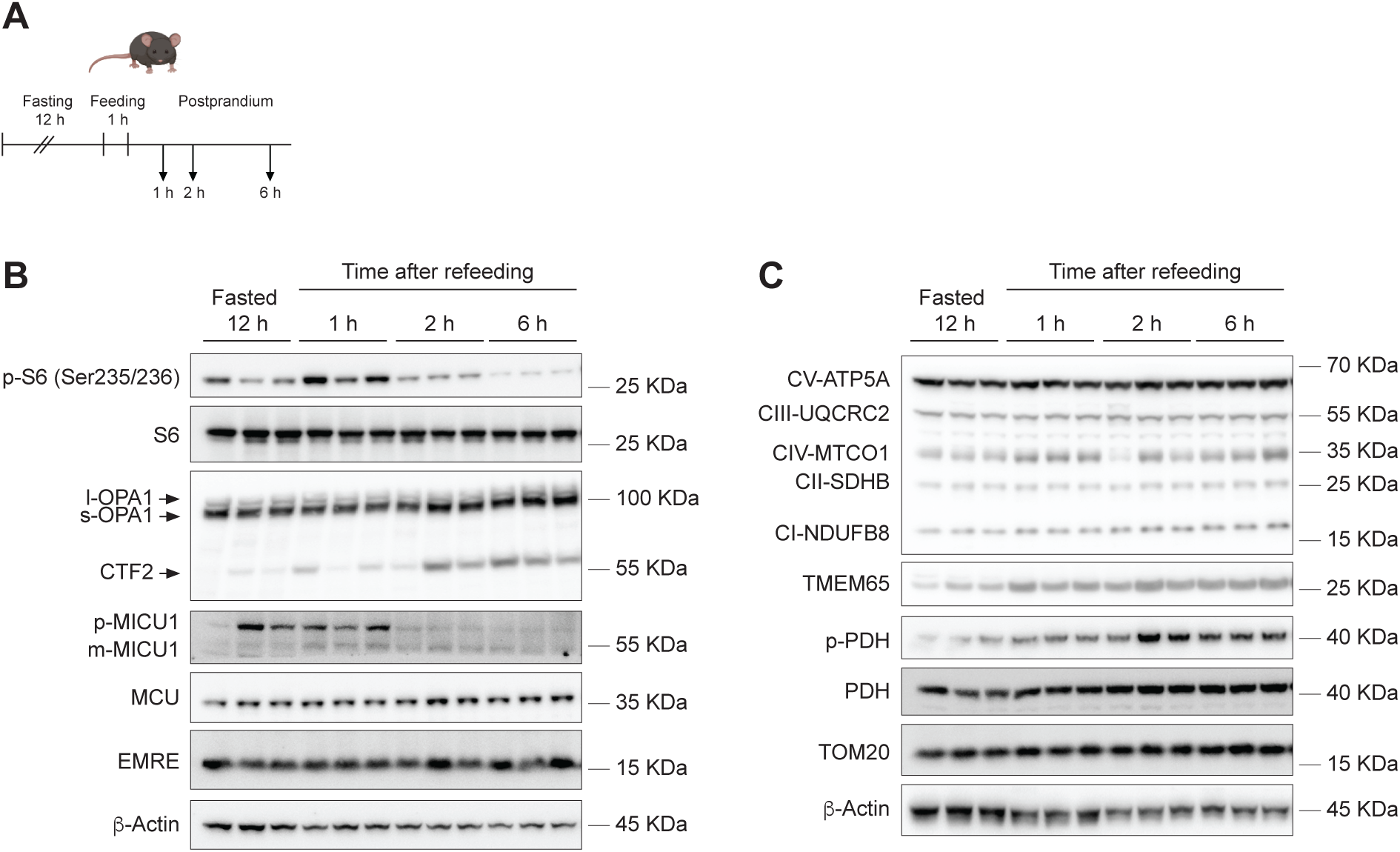
The proportion of non-gated mitochondrial Ca^2+^ uniporter complexes increases during fasting in postprandial mouse liver. Mice were fasted overnight for 12 hours and then refed for 1 hour. Livers were collected at 1, 2, and 6 hours after refeeding. (**A**) Experimental design. (**B**) and (**C**) Representative Western blots of whole-liver lysates collected as in (A) and probed with the indicated antibodies. β-actin was used as a loading control. n≥3.

We next evaluated the protein levels of the components of the MCUc. MCU levels increased slightly and consistently up to 6 hours after refeeding, as did EMRE, although the latter changes did not reach statistical significance (**Fig. 1B, S1D and S1E**). Notably, MCU/EMRE levels remained constant over time (**Fig. S1F**). The mature form of MICU1 (m-MICU1) transiently rose after refeeding but rapidly declined, reaching the lowest levels during fasting (**Fig. 1B and S1G**). In contrast, the precursor form of MICU1 (p-MICU1) markedly increased during fasting and in the early postprandial hours (**Fig. 1B**). The m-MICU1/MCU ratio, an indicator of the proportion of gated versus non-gated MCUc, followed a similar trend (**Fig. S1H**). MICU2 and MCUb levels were not assessed due to the lack of reliable antibodies for their immunodetection in this model. Together, these results suggest that the abundance of conductive, non-gated MCUc increases in the postprandial liver as a result of m-MICU1 downregulation.

Since MCU and EMRE levels showed a modest yet consistent postprandial increase, we next wondered whether this reflected specific MCUc remodelling or a broader enhancement of mitochondrial mass. To investigate this, we measured mitochondrial mass in whole-liver homogenates by Western blotting (**Fig. 1C**). Representative proteins that participate in oxidative phosphorylation, including UQCRC2 (Complex III), SDHB (Complex II), NDUFB8 (Complex I), and ATP5A (Complex V, ATP synthase) were upregulated following refeeding (**Fig. 1C and S1I-M**). In addition, the levels of the mitochondrial import receptor TOM20 and TMEM65, a protein involved in Na^+^-dependent Ca^2+^ efflux ^12, 13, 14^, also showed a trend toward increase (**Fig. 1C, S1N and S1O**). The ratio between the phosphorylated form (p-PDH) to non-phosphorylated PDH was significantly elevated (**Fig. 1C and S1P**). Collectively, these data indicate that postprandial conditions are accompanied by a delayed and global increase in mitochondrial mass, whereas an early and transient response to refeeding is the temporary accumulation of non-gated MCUc.

### MICU1 transcript and protein levels increase during serum deprivation and insulin signalling

Given the plasticity of mitochondrial Ca^2+^ uptake proteins during the transition between nutritional states *in vivo*, we next examined MICU1 levels *in vitro* in HeLa cells subjected to serum deprivation and insulin stimulation. Overnight foetal bovine serum (FBS) deprivation (12 hours; **Fig. 2A**) led to a marked increase in the precursor form of MICU1 (p-MICU1) (**Fig. 2B and S2A**). In contrast, the protein levels of m-MICU1, MCU, and MICU2 remained unchanged (**Fig. 2B and S2B-D)**. Subsequent incubation of serum-starved cells with 200 nM insulin enhanced p-MICU1 levels (**Fig. 2B and S2A**), while m-MICU1, MCU, and MICU2 levels remained unaffected (**Fig. 2B and S2B-D)**. Given the significant increase in p-MICU1 observed under both serum deprivation and insulin treatment, we next investigated whether these conditions also affected MICU1 mRNA expression. Consistent with the protein data, FBS deprivation specifically upregulated MICU1 mRNA, whereas insulin treatment did not further enhance its expression (**Fig. 2C**). Notably, MCU mRNA levels remained unchanged across conditions (**Fig. 2D**).

**Figure 2.**
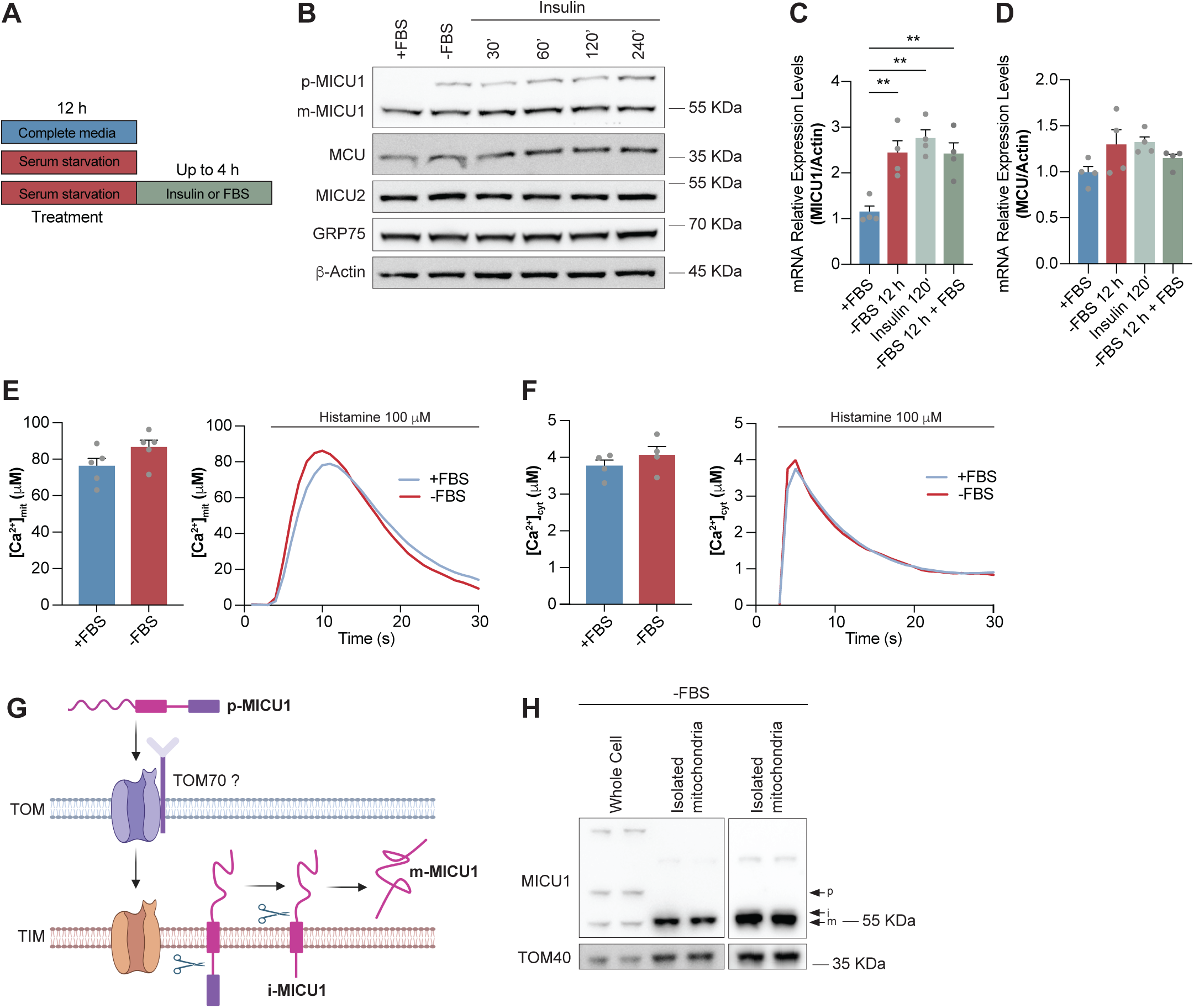
Serum deprivation increases p-MICU1 content, and insulin treatment further enhances p-MICU1 levels. (**A**) Schematic representation of the experimental protocol used for serum deprivation and insulin stimulation in HeLa cells. (**B**) Representative Western blot of HeLa cells cultured in complete medium (+FBS), deprived of FBS for 12 hours (-FBS), or deprived of FBS for 12 hours and the treated with 200 nM insulin for 30, 60, 120, or 240 minutes, probed with the indicated antibodies. GRP75 and β-actin were used as loading controls. (**C**) and (**D**) Quantitative PCR (qPCR) analysis of MCU and MICU1 mRNA expression levels in HeLa cells cultured in complete medium (+FBS), deprived of FBS for 12 hours (-FBS), and deprived of FBS for 12 hours and treated with 200 nM insulin for 120 minutes and FBS-deprived for 12 hours and reincubated with FBS for 4 hours. mRNA expression was normalized to actin. Data are presented as mean ± SEM. n=4. For data analysis, one-way ANOVA with post hoc Bonferroni tests was used. ** p≤0.01. (**E**) and (**F**) Mitochondrial ([Ca^2+^]_mit_) and cytosolic Ca^2+^ concentration ([Ca^2+^]_cyt_) measurements, respectively, in intact cells cultured in growth media or deprived of FBS for 12 hours and transfected with mitochondrial or cytosolic aequorin 48 hours before measurement. Cells were challenged with maximal histamine stimulation. Right panels: representative traces; left panels: bar diagram representing the mean Ca^2+^ peak upon stimulation ± SEM. n=5 for each condition for [Ca^2+^]_mit_ measurements; n=4 for each condition for [Ca^2+^]_cyt_ measurements. **(G)** Schematic representation of MICU1 processing and mitochondrial import (According to ^29^). **(H)** Whole-cell or mitochondria-enriched fractions obtained by differential centrifugation were analysed by Western blot for MICU1 precursor (p-MICU1), intermediate (i-MICU1), and mature (m-MICU1) forms. TOM40 was used to assess outer mitochondrial membrane integrity.

We then investigated whether the elevated p-MICU1 expression following FBS deprivation affected mitochondrial Ca²⁺ homeostasis. To this end, we measured both mitochondrial and cytosolic Ca^2+^ concentrations ([Ca^2+^]_mit_ and [Ca^2+^]_cyt_, respectively). Neither [Ca^2+^]_mit_ nor [Ca^2+^]_cyt_ were affected after 12 hours of serum deprivation (**Fig. 2E and 2F**), although mitochondrial Ca^2+^ tended toward an increase (**Fig. 2E**). Since p-MICU1 is not a part of the MCUc, we next investigated whether the MICU1 precursor accumulates within mitochondria or remains in the cytoplasm, potentially reflecting an impairment in mitochondrial protein import. Furthermore, MICU1 has been reported to associate with the MICOS complex, contributing to the regulation of cristae architecture ^32^. Thus, the increase in p-MICU1 might influence mitochondrial function through an MCU-independent mechanism. To explore this possibility, we isolated mitochondria-enriched fractions from cells deprived of FBS. Both the intermediate (i-MICU1) and mature (m-MICU1) forms were enriched in these mitochondrial fractions (**Fig. 2G and 2H**), whereas p-MICU1 was not detected (**Fig. 2H**). These results indicate that, although p-MICU1 accumulates under FBS deprivation, its processing into mitochondrial forms is not proportionally increased. This suggests altered MICU1 maturation, potentially reflecting impaired import or processing efficiency.

Finally, to further explore models of altered nutrient sensing, we inhibited mTORC1 with rapamycin in HeLa cells (**Fig. S3**). Rapamycin treatment induced the expected dephosphorylation of S6 and increased phosphorylation of AKT (**Fig. S3A**). In contrast, MCU, MICU2 and m-MICU2 protein levels were unaffected (**Fig. S3A-D**), while p-MICU1 levels decreased (**Fig. S3A and S3E**). Overall, these findings indicate that diverse changes in nutrient availability and signalling converge to modulate MICU1 expression and processing.

### Glutamine deprivation increases mitochondrial Ca^2+^ uptake and non-gated MCUc

Glutamine is an essential metabolite in nutrient sensing, regulating multiple aspects of cellular function through the nutrient sensor mTORC1 ^33^. Although mammalian cells can synthesize glutamine, it may become limiting under certain metabolic conditions ^34, 35^. To investigate how cells adapt to glutamine deprivation, we explored cell adaptations to glutamine deprivation focusing on mitochondrial Ca^2+^ transport and metabolism. In HeLa cells, glutamine deprivation significantly reduced the levels of p-MICU1, m-MICU1, and MICU2 without changes in MCU (**Fig. 3A-E**). Of note, treatment of these cells with mTOR inhibitor rapamycin also modulated p-MICU1 levels, without changes in other components of the MCUc complex (**Fig. S3**). Consistently, HEK-293 cells also displayed decreased p-MICU1 upon glutamine deprivation (**Fig. S4A and S1B**), indicating that MICU1 levels are sensitive to nutrient availability across different cell types. In contrast, mature MICU1, MCU, and EMRE expression remained unchanged (**Fig. S4A and S4C-E**). In HEK-293 cells, the loss of p-MICU1 caused by glutamine starvation, but not that of mature MICU1 and MCU, could be prevented by the proteasome inhibitor MG132 (**Fig. S4F-I**), suggesting that it occurs through protein degradation. Unexpectedly, EMRE expression was blunted by MG132 (**Fig. S4F and S4J**).

**Figure 3.**
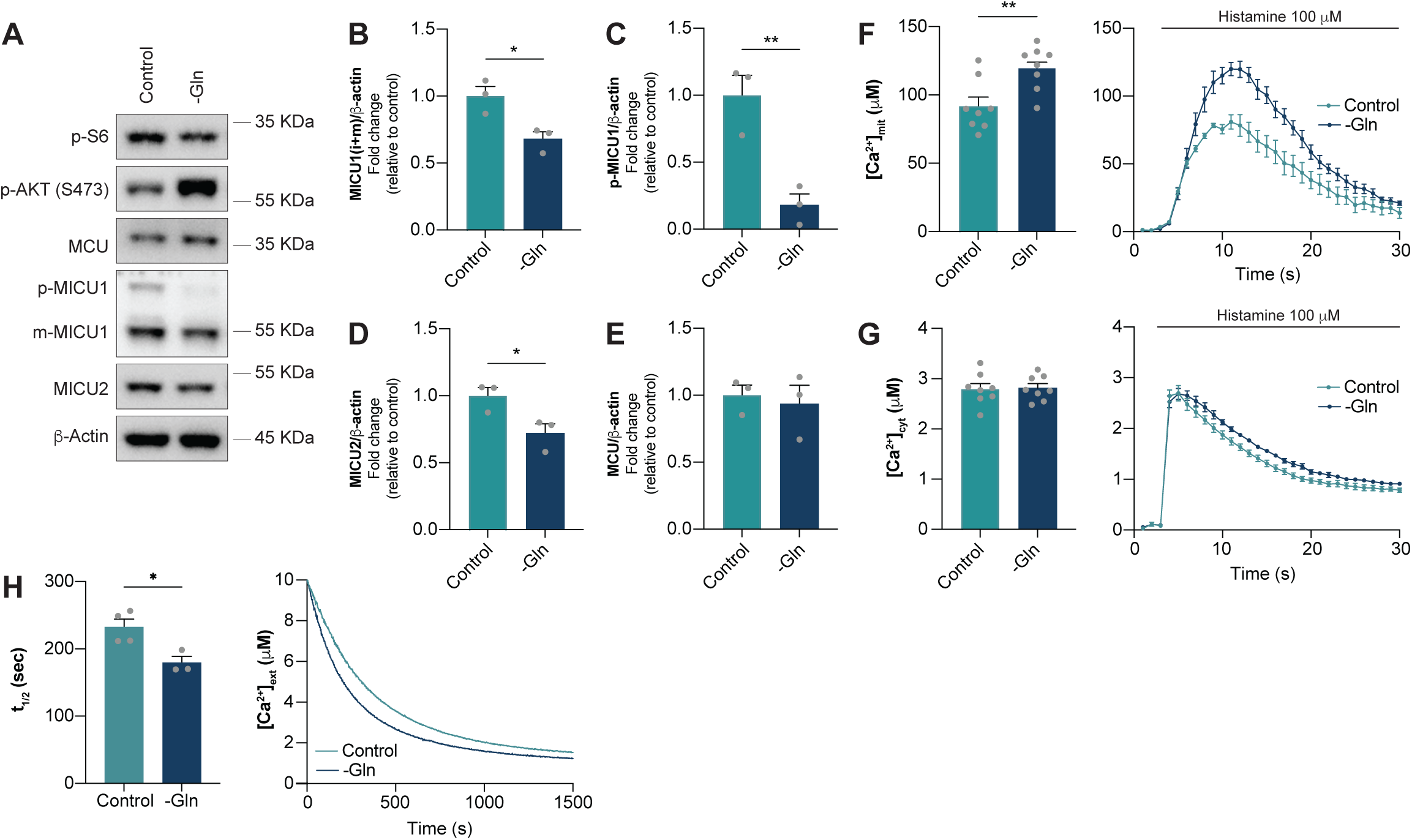
Glutamine deprivation induces MCUc remodelling and increases mitochondrial Ca^2+^ uptake. (**A**) Representative Western blots of HeLa cells cultured in complete media (Control) or glutamine-deprived media (-Gln) for 16 hours probed with the indicated antibodies. β-actin was used as loading control. (**B**) - (**E**) Quantification of MCUc components from the Western blots shown in (A). Data are presented as mean ± SEM. n=3. For analysis, unpaired Student T-test was used. * p≤0.05; ** p≤0.01. (**F**) and (**G**) [Ca^2+^]_mit_ and [Ca^2+^]_cyt_ measurements in intact cells cultured in complete media (Control) and glutamine-deprived media (-Gln) and transfected with mitochondrial and cytosolic aequorin 48 hours before Ca^2+^ measurements. Cells were challenged with maximal histamine stimulation. Right panels: representative traces; left panels: bar diagram representing the mean Ca^2+^ peak upon stimulation ± SEM. n=8. For analysis, unpaired Student T-test was used. * p≤0.05; ** p≤0.01. (**H**) Mitochondrial Ca^2+^ uptake in digitonin-permeabilized cells. The decrease in extramitochondrial Ca^2+^ levels ([Ca^2+^]_ext_), was monitored using Calcium Green-5N. Data were fitted to a one-phase decay equation, and uptake rates were derived from the half-life values. Data are presented as mean ± SEM. n≥3. For analysis, unpaired Student T-test was used. * p≤0.05.

To assess the functional consequences of these alterations in MCUc composition, the genetically-encoded sensor aequorin was used to measure [Ca^2+^]_mit_ and [Ca^2+^]_cyt_ in cells deprived of glutamine overnight ^36^. Histamine was added to trigger Ca^2+^ transients in both compartments (**Fig. 3F, 3G**). Mitochondrial Ca^2+^ uptake was markedly enhanced in response to glutamine removal (**Fig. 3F**), while cytoplasmic Ca^2+^ signals remained unchanged (**Fig. 3G**). Mitochondrial steady state [Ca^2+^] reflects a dynamic equilibrium between Ca^2+^ uptake, Ca^2+^ buffering, and Ca^2+^ efflux. Therefore, the observed increase in mitochondrial Ca^2+^ uptake may result from greater Ca^2+^ transfer from the endoplasmic reticulum (ER), higher mitochondrial membrane potentials (ΔΨ), or increased inner mitochondrial membrane (IMM) permeability to Ca^2+^, possibilities we investigated next.

Co-localization analyses using TOM20 to label mitochondria and SEC61-β-GFP to label the ER revealed no detectable changes in ER-mitochondria contact sites (**Fig. S5A**). Mitochondria from cells under glutamine starvation appear more elongated, thin, and connected (**Fig. S5B**), which is in accordance to previous data ^37^. An elongated mitochondrial phenotype has been associated to an increased ability to uptake Ca^2+^, a complex and not yet well understood phenomena ^38^. To confirm that the enhanced mitochondrial Ca^2+^ uptake in response to glutamine deprivation was independent of ER-mitochondrial Ca^2+^ exchange, we measured mitochondrial Ca^2+^ uptake in digitonin-permeabilized cells ^26, 38, 39^, a condition that disrupts cytosolic Ca^2+^ microdomains. ATP was added in the assay media to inhibit mitochondrial permeability transition ^40^, allowing the determination of maximal Ca^2+^ uptake rates. We found that mitochondria from glutamine-deprived cells take up Ca^2+^ at higher rates, even in the absence of ER-derived Ca^2+^ release (**Fig. 3H**). These results are consistent with an MICU1-deficient MCUc configuration ^41^, leading to enhanced mitochondrial ion permeability and elevated [Ca^2+^]_mit_ measured in intact cells. Collectively, our findings show that glutamine deprivation promotes the loss of MICU1 and MICU2, resulting in a non-gated MCUc and exacerbated mitochondrial Ca^2+^ uptake.

### Glutamine deprivation increases sensitivity to mitochondrial permeability transition

Ca^2+^ regulates substrate transport and oxidation in mitochondria, thus enhancing oxygen consumption rates (OCR) within a specific concentration range ^4, 42^. As MICU1 loss is associated with higher mitochondrial Ca^2+^ levels ^41^, we hypothesized that mitochondrial OCR might be upregulated under glutamine deprivation. To test this, we measured OCR and extracellular acidification rates (ECAR) in viable cells using an extracellular flux analyser (**Figure 4A-C**). Briefly, cells were cultured overnight in complete medium or under glutamine-free conditions, and OCR/ECAR were measured in both groups under basal conditions. Glutamine was then added acutely (to reconstitute complete medium), followed by sequential addition of oligomycin (to inhibit ATP synthase) and antimycin A plus rotenone (to determine mitochondrial OCR). Acute glutamine addition promoted a larger increase in OCR in glutamine-starved cells, suggesting a higher capacity to oxidize glutamine (**Fig. 4A and 4B**). The difference in basal OCR (after glutamine readdiction) between starved cells and controls is likely a consequence of higher ATP production, as revealed by higher ATP-linked respiration (**Figure 4B**). A small non-significant difference in OCR persisted after oligomycin addition (**Fig. 4B**). This slightly higher oligomycin-sensitive respiration observed in the glutamine-starved cells is likely related to increased ion transport rates, which dissipate the electrochemical proton gradient independently of ATP synthase. No significant differences were detected in non-mitochondrial respiration (**Fig. 4B)** or in ECAR values between the groups (**Fig. 4C**).

**Figure 4.**
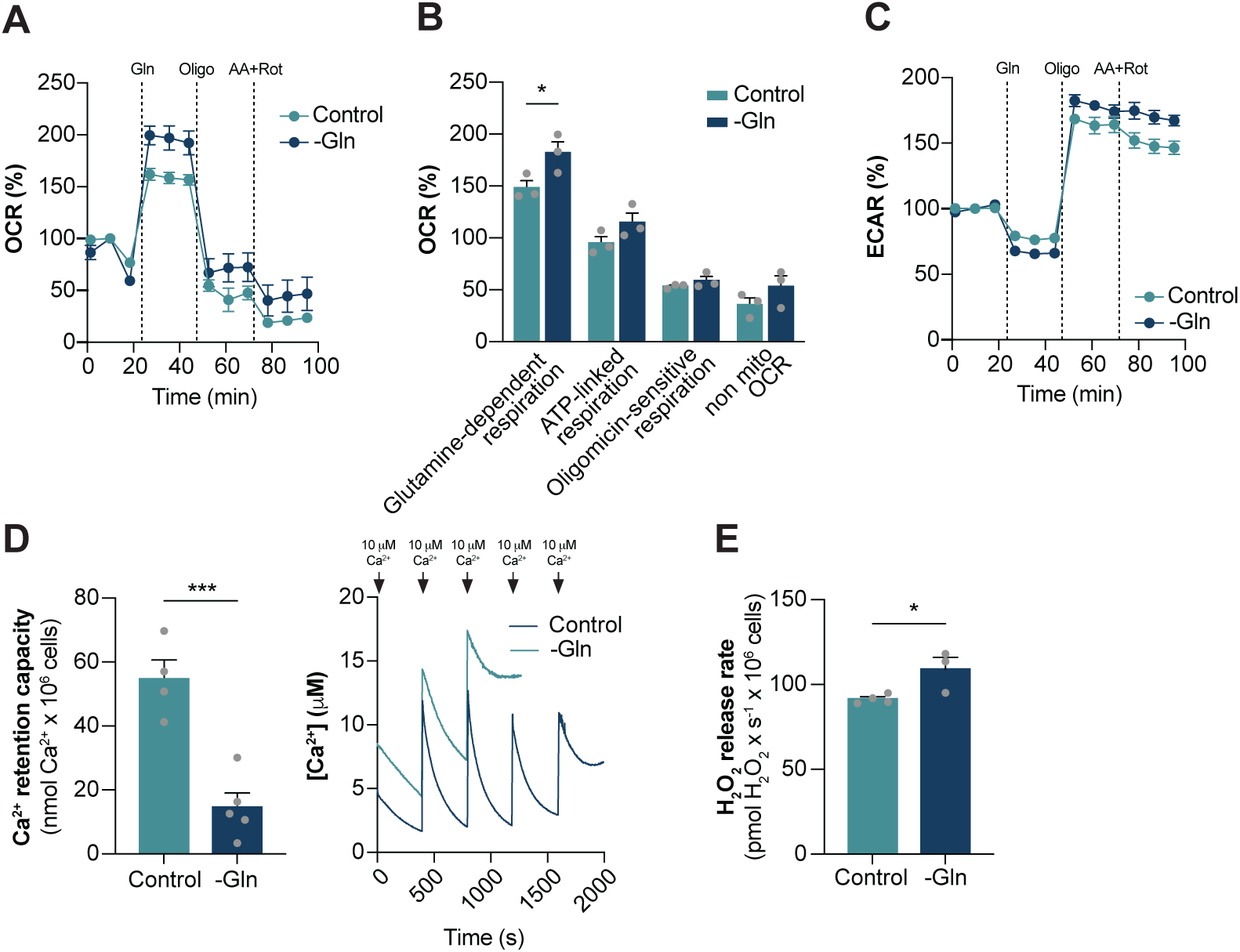
Glutamine deprivation alters mitochondrial oxygen consumption, Ca^2+^ retention capacity, and H_2_O_2_ release. Oxygen consumption rates (OCR) and extracellular acidification rates (ECAR) were measured in HeLa cells using an extracellular flux analyser (Seahorse XFe24). Cells were cultured overnight in complete medium (Control) or glutamine-free medium (-Gln). **(A)** Representative plots of OCR. Baseline measurements were obtained in DMEM media without glutamine, for both groups, followed by sequential additions of 4 mM glutamine, 1 µM oligomycin, and 1 µM rotenone plus 1 µM antimycin A.Data were normalized to the baseline measurements. **(B)** Glutamine-dependent respiration, ATP-linked respiration, oligomycin-sensitive respiration and non-mitochondrial respiration were calculated from the experiment shown in A as previously performed ^52^. Data are presented as mean ± SEM. n=3. For analysis, unpaired Student T-test was used. * p≤0.05. **(C)** Representative plots of ECAR. Baseline measurements were obtained in DMEM media without glutamine, for both groups, followed by sequential additions of 4 mM glutamine, 1 µM oligomycin, and 1 µM rotenone plus 1 µM antimycin A. Data were normalized to the baseline measurements. **(D)** Mitochondrial Ca^2+^ uptake measurements in digitonin-permeabilized HeLa cells grown in complete media (Control) or in the absence of glutamine (-Gln), monitored with Calcium Green-5N. Successive 10 µM Ca^2+^ additions were performed until permeability transition occurred, indicated by a sharp fluorescence increase. Right panel: representative traces of the experiment. Left panel: quantification of maximal Ca²⁺ retention capacity. Data are presented as mean ± SEM. n=5. For analysis, unpaired Student T-test was used. *** p≤0.001. **(E)** Mitochondrial H_2_O_2_ release in permeabilized HeLa cells. Data represent as mean H_2_O_2_ release, normalized by number of cels ± SEM. For analysis, unpaired Student T-test was used. *p≤0.05.

We next quantitatively measured the capacity of mitochondria to take up and store Ca^2+^ in permeabilized cells using the extramitochondrial sensor Ca^2+^ Green-5N. Repeated Ca^2+^ boluses were added to the suspension, and fluorescence was monitored as mitochondria actively took up Ca^2+^, leading to a decrease in extramitochondrial fluorescence, until the induction of permeability transition, which causes fluorescence to rise again. Mitochondria from glutamine-starved cells exhibited an increased propensity for permeability transition and thus display a reduced Ca^2+^ retention capacity (**Fig. 4D**). Mitochondrial permeability transition is typically associated with redox imbalance, with enhanced mitochondrial reactive oxygen species (ROS) formation, triggered by excessive mitochondrial Ca^2+^ uptake ^40^. Consistent with this, we found elevated mitochondrial hydrogen peroxide (H_2_O_2_) release in glutamine-deprived cells, as measured by Amplex Red oxidation (**Fig. 4E**).

### YME1L is necessary for MICU1 degradation

We next investigated the mechanisms by which nutrient availability regulates MICU1 stability. Initially, we focussed on mitochondrial proteases OMA1, AFG3L2, and YME1L, which are involved in the processing of several IMM and intermembrane space (IMS) mitochondrial proteins [reviewed by ^43^].

OMA1 was activated by inducing membrane depolarization with CCCP ^44^. This treatment did not affect mature MICU1 protein levels (**Fig. S6A and S6B**), although OMA1 activation was induced, as assessed through the levels of l-OPA1 and s-OPA1 [**Fig. S6A** and ^44^]; interestingly, p-MICU1 and i-MICU1 decreased in response to mitochondrial depolarization (**Fig. S6A, S6C and S6D**), suggesting an increase in turnover by OMA1 and/or import. Conversely, OMA1 silencing decreased MICU1 levels under basal conditions (**Fig. S6E and S6F**). Furthermore, silencing of AFG3L2 produced approximately a two-fold increase in TMEM65 expression (**Fig. S6E and S6G**). Altogether, these results suggest that neither OMA1 nor AFG3L2 play a major role in MICU1 degradation.

In contrast, YME1L silencing led to m-MICU1 upregulation under basal conditions (**Fig. 5A and B**). This effect is specific for MICU1, since TMEM65, EMRE, MCU, and ATP5A are not affect by YME1L silencing (**Fig. 5C-F**). Moreover, YME1L silencing prevented glutamine deprivation-stimulated MICU1 loss and abolished the associated increase in mitochondrial Ca^2+^ uptake (**Fig. 5H-I**). In cells transfected with non-targeting RNA sequence, and under glutamine starvation, a minute and constant decrease in cytosolic Ca^2+^ peak was observed, probably as a consequence of the sequence of stress triggering events. Nevertheless, the increase in mitochondrial Ca^2+^ uptake remained after glutamine deprivation and corrected by YME1L silencing (**Fig. 5H-I**). Altogether, these findings identify YME1L as a key regulator of MICU1 turnover and suggest that it functions as a nutrient-sensitive mitochondrial protease mediating MCUc remodelling in response to metabolic cues.

**Figure 5.**
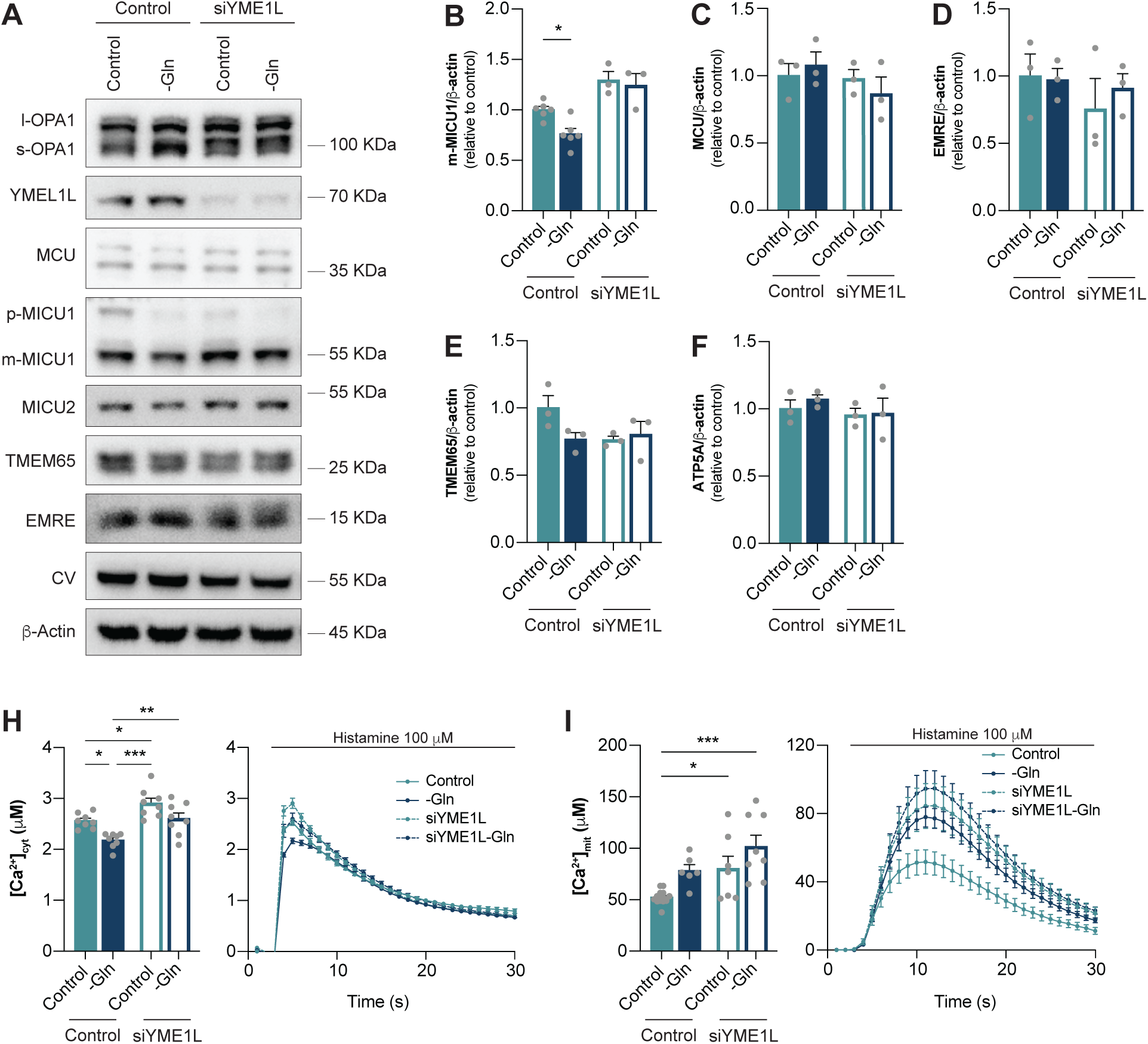
YME1L is required for MICU1 degradation induced by glutamine deprivation in HeLa cells. HeLa cells were transfected with control or YME1L siRNA for 48 hours and then incubated in complete media (Control) or media without glutamine (-Gln). (**A**) Representative Western blot probed with the indicated antibodies. β-actin was used as loading control. (**B**) - (**G**) Densitometric analyses for m-MICU1 (B), MCU (C), EMRE (D), TMEM65 (E), and ATP5A (F), normalized to β-actin. Data are presented as fold change relative to Control and represent the mean ± SEM. n≥3. For data analysis, one-way ANOVA with post hoc Bonferroni tests was used. * p≤0.05. (**H**) - (**I**) Cytosolic ([Ca^2+^]_cyt_) and mitochondrial (Ca^2+^]_mit_) Ca^2+^ levels, respectively, measured using aequorin-based probes after maximal histamine stimulation. n=8. Data are shown as mean ± SEM. For data analysis, one-way ANOVA with post hoc Bonferroni tests was used. *p≤0.05; ** p≤0.01; *** p≤0.001.

## Discussion

Here we demonstrate that MCUc components and mitochondrial Ca^2+^ uptake are highly responsive to nutrient availability. We previously showed that long-term caloric restriction *in vivo* increases mitochondrial Ca^2+^ uptake rates in mitochondria from brain, liver, and kidney ^8, 9, 10^. Similarly, we now show that glutamine deprivation *in vitro* increases mitochondrial Ca^2+^ uptake rates, providing further evidence of the high plasticity of the mitochondrial Ca^2+^ transport machinery in response to nutrient status. These observations are consistent with the findings by Marchi and co-workers ^29^, who reported that glucose starvation enhances mitochondrial Ca^2+^ uptake via MICU1 downregulation. Indeed, we also observed a decrease in MICU1 under glutamine deprivation *in vitro*.

Additionally, our *in vivo* fasting-refeeding experiments further revealed that MICU1 levels change dynamically over short time scales: they are low during fasting, increase rapidly upon refeeding, and subsequently decline within a few hours. This highlights that MCUc plasticity, particularly involving MICU1 levels, acts as an acute and reversible mechanism of adaptation to nutrient fluctuations.

Nutrient status is known to modulate multiple mitochondrial components. In our fasting-refeeding model, markers of mitochondrial mass, including several electron transport chain components, increased progressively during the first 6 hours of refeeding. MCU and EMRE, two components of the MCUc, followed a similar trend. Levels of TMEM65, a protein involved in mitochondrial Ca^2+^ efflux ^12, 13, 14^, were reduced during starvation and gradually increased upon refeeding. In contrast, MICU1 levels exhibited a rapid and transient rise followed by a sharp decline, leading to a lower m-MICU1/MCU ratio between 2-6 hours post-refeeding. This pattern suggests a transient accumulation of conductive, non-gated MCUc ^45^, consistent with enhanced mitochondrial Ca^2+^ uptake.

In glutamine starved cells, the loss of MICU1 was followed by a similar decrease in MICU2 levels (not quantified *in vitro* due to antibody limitations). We hypothesize that low MICU2 levels occur secondarily to m-MICU1 downregulation of, since loss of m-MICU1 favours MICU2 degradation. Indeed, previous studies showed that MICU1/MICU2 or MICU1/MICU1 dimers are more resistant to proteolytic degradation than monomeric forms ^46^.

To determine whether the higher proportion of non-gated, conductive MCUc generated in response to nutritional status had functional consequences, we measured mitochondrial Ca^2+^ concentrations. In intact cells subjected to glutamine deprivation, histamine-induced mitochondrial Ca^2+^ uptake was enhanced. These results contrast with some MICU1 knockdown and knockout models reporting reduced mitochondrial Ca^2+^ uptake, a discrepancy that may arise from concurrent downregulation in those systems, which produces non-gated yet non-conductive MCUc channels ^46^. In our model, EMRE levels remained stable after nutrient deprivation, and only m-MICU1/MCU ratio was altered.

Direct measurements of mitochondrial Ca^2+^ uptake in permeabilized cells, where plasma membrane and ER Ca^2+^ fluxes are excluded ^38^, confirmed that nutrient deprivation directly enhances mitochondrial Ca^2+^ transport. Measurements of dynamic Ca^2+^ uptake rates in permeabilized cells required the presence of ATP in the experimental media to prevent Ca^2+^-induced permeability transition. Consistently, glutamine-deprived cells exhibited lower Ca^2+^ retention capacity and higher release of H_2_O_2_, indicating higher susceptibility to permeability transition. These results closely mirror our previous observations in caloric restriction models in kidney, where mitochondria show faster Ca^2+^ uptake, greater H_2_O_2_ generation, and increased susceptibility to the permeability transition ^10^.

Glutamine is particularly important to support energetic metabolism boost under stress ^47^. Its metabolism through glutaminase and/or asparagine synthetase provides substrates such as αketoglutarate that can be readily oxidized by mitochondria. Increased mitochondrial Ca^2+^ uptake is known to enhance electron transport rates, TCA cycle flux, oxidative phosphorylation, and ROS release ^5, 42^. Accordingly, cells deprived of glutamine displayed higher basal oxygen consumption rates (OCR) (after glutamine replenishment) and increased ROS release compared to control cells. We propose that the increase in mitochondrial Ca^2+^ that occurs as consequence of MCUc remodelling stimulates TCA cycle Ca^2+^-sensitive enzymes (or substrate transporters), such as α-ketoglutarate dehydrogenase, increasing OCR and ROS production.

We next sought to identify the mechanism underlying m-MICU1 downregulation in response to changes in nutrient availability, hypothesizing a protease-dependent process. Through a targeted screening approach, we identified the i-AAA protease YME1L as responsible for nutrient-sensitive degradation of MICU1. YME1L silencing increased m-MICU1 levels under basal conditions and prevented its downregulation during glutamine deprivation, without altering MCU, EMRE, nor mitochondrial mass markers. These experiments demonstrate that nutrient-dependent regulation of MICU1 is mediated by YME1L. Furthermore, TMEM65 downregulation during glutamine deprivation was also prevented by YME1L silencing. Consistently, YME1L is an IMM protease whose proteolytic domain faces the IMS ^48^. Supporting this mechanism, studies in YME1L knockout HEK-293 cells exposed to hypoxia, conditions that modulate mTOR signalling, identifies MICU1 as a potential YME1L substrate ^35^.

Overall, our findings reveal that mitochondrial Ca^2+^ transport is dynamically and specifically regulated by nutritional status. MICU1 turnover, mediated by the mitochondrial protease YME1L, represents a key mechanism through which mitochondrial Ca^2+^ uptake and MCUc composition are remodelled during fasting/feeding transitions, nutrient deprivation, and hormonal signalling. This identifies YME1L and MICU1 as central players in the metabolic remodelling of mitochondria in response to nutrient availability.

## Materials and Methods

### Postprandial protocol: fasting/feeding transition

Male C57BL/6NCrl mice (Charles River), 8-12 weeks old, were lodged at the Animal facility of the Biomedical Sciences Department, University of Padova. Experiments were approved by the local ethics committee in accordance with Italian law D.L.vo n° 26/2014. Animals were maintained under specific pathogen–free conditions at constant temperature and a 12 h light/12 h dark photoperiod (lights on at 6:00). Mice were fasted overnight (7:00 PM-7:00 AM) with free access to water (Control). In the morning, they were allowed to feed for 1 hour and subsequently transferred into clean cages. Animals were euthanized at the indicated time points (fasted, or 1, 2, or 6 hours after refeeding).

### Cell cultures

HeLa or HEK-293 cells were grown in high-glucose DMEM (DMEM-HG: 1 mM pyruvate and 4 mM glutamine) supplemented with 10% FBS and 1% penicillin/streptomycin. After 24 hours, cells were washed three times with PBS (or DPBS in the case of HEK-293 cells) and incubated with complete DMEM-HG (Control) or in media lacking specific nutrients or components, as described in figure legends.

### Serum starvation and insulin stimulation

HeLa cells were seeded in complete DMEM-HG and allowed to grow for 24 hours. Cells were then washed three times with PBS and incubated for 16 hours in FBS-free media. After this period, media was replaced to DMEM-HG containing penicillin/streptomycin and either water (Control) or insulin (200 nM). For serum replenishment experiments, media was replaced to complete DMEM-HG supplemented with FBS (or not, Control).

### Glutamine starvation and rapamycin treatment

HeLa or HEK-293 cells were seeded and grown for 24 hours in DMEM-HG and 10% FBS to reach 70-80% confluence. Cells were then washed three times with PBS (or Dulbeccós D-PBS for HEK-293 cells) and incubated for 16 hours with DMEM-HG supplemented with 5% FBS, 1 mM pyruvate, and 4 mM glutamine (Control), or the same media without glutamine. For rapamycin experiments, HeLa or HEK-293 cells were grown to 70–80% confluence, then switched to DMEM-HG with 10% FBS supplemented with 1 mM pyruvate and 4 mM glutamine, in the presence or absence of 200 nM rapamycin for 16 hours. Cells were subsequently collected for analysis.

### Western blots

Cells or mitochondria-enriched fractions were lysed in RIPA buffer (150 mM NaCl, 50 mM Tris-HCl pH=8, 1 mM EGTA, 1% Triton X-100, 0.5% sodium deoxycholate, and 0.1% SDS), supplemented with Complete EDTA-free protease inhibitor cocktail (Roche Applied Science), and Phosphatase inhibitor cocktail (Roche Applied Science). Liver samples were homogenized in T-PER (Thermo Fisher Scientific, #78510). Protein concentrations were measured by bicinchoninic acid assay. Samples were denatured in Laemmli buffer containing 1 mM dithiothreitol (DTT) at 95° C for 5 minutes. Equal amounts of proteins (25-40 µg) were resolved by electrophoresis on Bis-Tris 4-12% precast gels (Thermo Fisher Scientific) and transferred onto 0.45 µm nitrocellulose membranes. Membranes were blocked in 5% non-fat milk dissolved in TBST (20 mM Tris, 150 mM NaCl, and 0.1% Tween 20; pH 7.4). Primary antibodies (Table 1) were diluted in 5% milk and incubated overnight. Membranes were incubated with the appropriate horseradish peroxidase (HRP)-conjugated secondary antibody. Images were analysed with the Fiji distribution of ImageJ. All figures represent at least three independent experiments.

**Table 1.**
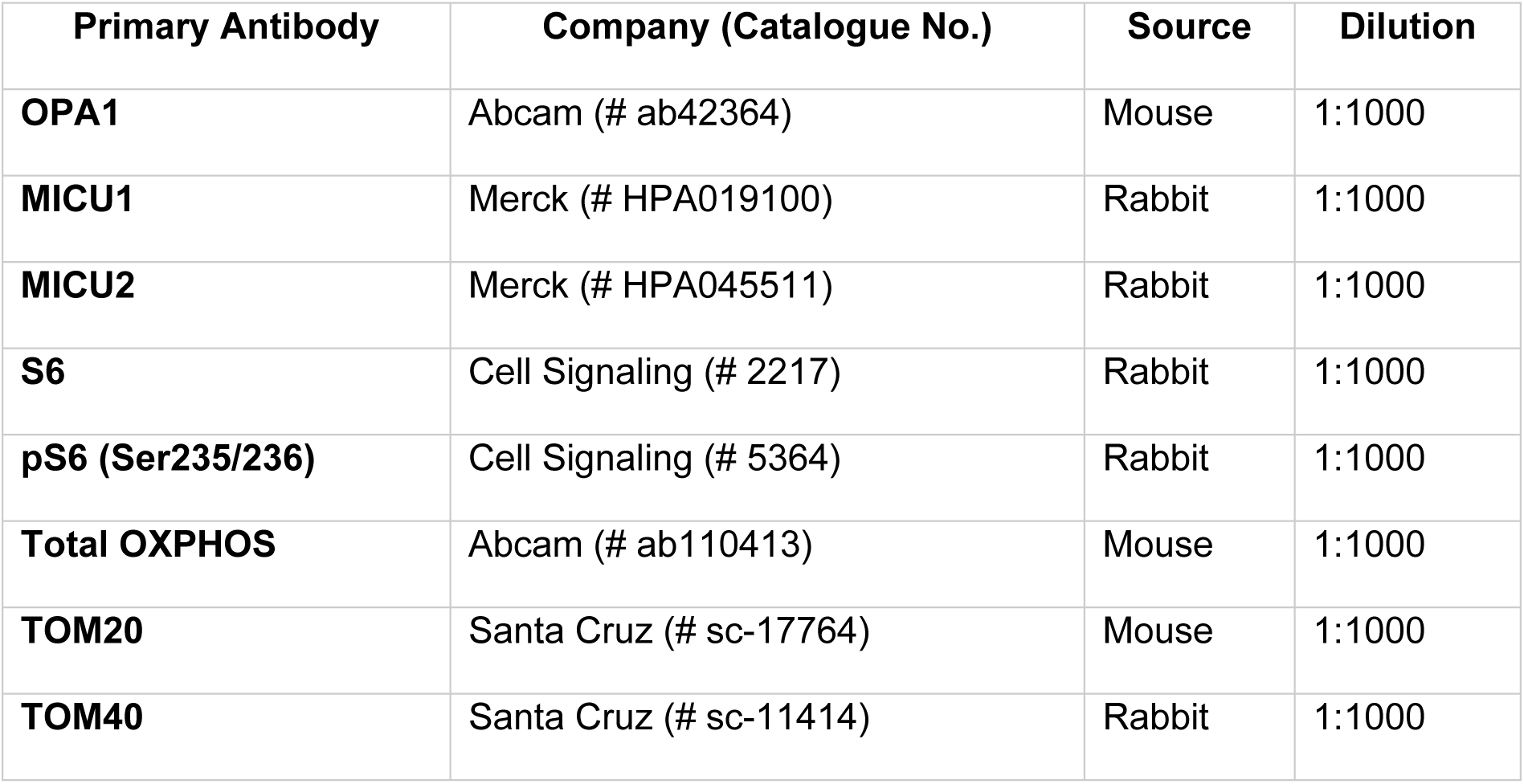
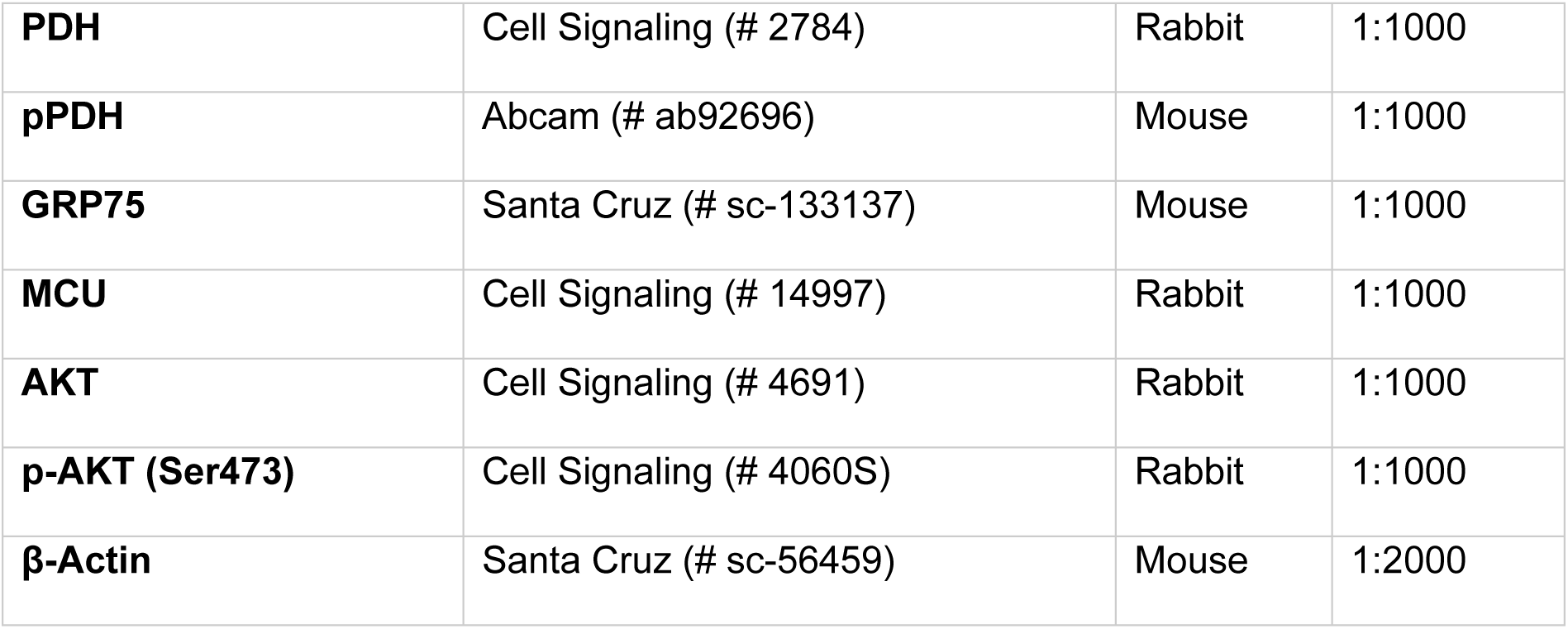
Primary antibodies.

| Primary Antibody | Company (Catalogue No.) | Source | Dilution |
| --- | --- | --- | --- |
| **OPA1** | Abcam (# ab42364) | Mouse | 1:1000 |
| **MICU1** | Merck (# HPA019100) | Rabbit | 1:1000 |
| **MICU2** | Merck (# HPA045511) | Rabbit | 1:1000 |
| **S6** | Cell Signaling (# 2217) | Rabbit | 1:1000 |
| **pS6 (Ser235/236)** | Cell Signaling (# 5364) | Rabbit | 1:1000 |
| **Total OXPHOS** | Abcam (# ab110413) | Mouse | 1:1000 |
| **TOM20** | Santa Cruz (# sc-17764) | Mouse | 1:1000 |
| **TOM40** | Santa Cruz (# sc-11414) | Rabbit | 1:1000 |
| **PDH** | Cell Signaling (# 2784) | Rabbit | 1:1000 |
| **pPDH** | Abcam (# ab92696) | Mouse | 1:1000 |
| **GRP75** | Santa Cruz (# sc-133137) | Mouse | 1:1000 |
| **MCU** | Cell Signaling (# 14997) | Rabbit | 1:1000 |
| **AKT** | Cell Signaling (# 4691) | Rabbit | 1:1000 |
| **p-AKT (Ser473)** | Cell Signaling (# 4060S) | Rabbit | 1:1000 |
| **β-Actin** | Santa Cruz (# sc-56459) | Mouse | 1:2000 |

### Cell permeabilization: digitonin titration

Digitonin was used to selectively permeabilize the plasma while preserving mitochondrial integrity ^39^. Optimal digitonin concentrations were determined using an O2k high-resolution oxygraphy (Oroboros Instruments), as previously described ^10^. A total of 10^6^ cells were resuspended in 2 ml of experimental buffer (0.3 M sucrose, 10 mM HEPES, 2 mM EGTA, 1 mM EDTA; pH 7.2) with 1 mM EGTA, 1 mM succinate, 1 mM rotenone and 1 mM ADP (state 3). Because intact cells poorly permeate succinate and ADP, oxygen consumption rates (OCR) remain low until permeabilization occurs. Digitonin was added in boluses until maximal, the restrains imposed by the plasma membrane are removed and respiration rates increase. The minimal digitonin concentration that produced maximal respiration was used for all experiments.

### Ca^2+^ transport assays

10^6^ cells were resuspended in 2 ml of experimental buffer (0.3 M sucrose, 10 mM HEPES, 2 mM EGTA, 1 mM EDTA; pH 7.2) containing 0.1 µM Calcium Green.5N, 1 mM succinate, and 1 µM rotenone. Fluorescence (excitation 506 nm; emission 532 nm) was recorded at 37° C with constant stirring using an F4500 Hitachi fluorimeter. Extramitochondrial Ca^2+^ concentrations were calculated using:

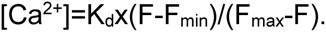

K_d_ was empirically determined from fluorescence changes elicited by a known 10 µM Ca^2+^ addition. Maximal (F_max_) and minimal (F_min_) fluorescence values were obtained at the end of each experiment by addition of 100 µL of 100 mM CaCl_2_ and 100 mM EGTA.

For uptake-rate measurements, 1 mM ATP was included to prevent permeability transition. After Ca^2+^ addition, fluorescence was allowed to decrease to a plateau. Data were fitted to a one-phase decay function, and half-lives were used to quantify mitochondrial Ca^2+^ uptake rates.

### Mitochondria isolation

Mitochondria-enriched fractions from HeLa cells were prepared through differential centrifugation as described by Frezza and co-workers ^49^ with modifications. Briefly, cells (around 50 x 10^6^ cells) were washed with PBS, starved of FBS overnight, trypsinized, and homogenized using first a Potter-Elvehjem and then a Dounce homogenizer. Samples were centrifuged at 200 x g, and the pellet, containing non-homogenized cells, was resuspended and re-homogenized as previously described. The supernatant was centrifuged at 900 x g to remove nucleus and cell debris. The supernatant was centrifuged at 9000 x g to pellet mitochondria, which were washed, resuspended in isolation buffer and pelleted again. Finally, the pellets were lysed in RIPA buffer (150 mM NaCl, 50 mM Tris-HCl pH=8, 1 mM EGTA, 1% Triton X-100, 0.5% sodium deoxycholate, and 0.1% SDS), supplemented with Complete EDTA-free protease inhibitor cocktail (Roche Applied Science), and Phosphatase inhibitor cocktail (Roche Applied Science) and denatured in Laemmli buffer for Western blotting.

### Aequorin-based measurements of mitochondrial and cytoplasmic Ca^2+^

Aequorin measurements were performed using a PerkinElmer EnVision plate reader equipped with a two-injector unit. Cells were transfected with aequorin plasmids with a standard Ca^2+^-phosphate method, as already performed ^50^. Aequorin was reconstituted by incubating cells for 1 hour with Krebs-Ringer buffer (KRB: 125 mM NaCl, 5 mM KCl, 1 mM Na_3_PO_4_, 1 mM MgSO_4_, 5.5 mM glucose, 20 mM HEPES; pH 7.4), supplemented with 5 µM coelenterazine. Luminescence was recorded for 1 minute, followed by injection of 100 μM histamine to evoke Ca²⁺ transients. At the end of each measurement, a hypotonic Ca²⁺-rich solution (10 mM Ca²⁺ + 100 μM digitonin) was added for calibration. Data were analysed as described by Pinton and collaborators ^36^, using a custom made macro-enabled Excel workbook.

### Mitochondrial morphology and co-localization experiments

HeLa cells (5 x 10^5^) were seeded on glass coverslips in complete DMEM-HG and allowed to grow for 24 hours. For ER-mitochondria co-localization experiments, cells were transfected with 0.2 μg Sec61- β-GFP plasmid (gift from Prof. Paola Pizzo) using the standard Lipofectamine 3000 reagent protocol (Thermo Fisher, #L3000001) in FBS-free media for 4 hours, after which growth media was restored. Cells were subjected to glutamine starvation, washed with PBS and fixed with 4% PFA at 4° C for 20 minutes.

Fixed cells were washed, quenched with 50 mM NH_4_Cl for 20 minutes, permeabilized with 0.1% Triton X-100 for 5 minutes, and blocked with 3% FBS + 1% normal goat serum for 1 hour. Cells were incubated overnight at 4° C with an anti-TOM20 (1:400), followed by Alexa Fluor 594–conjugated secondary antibody (1:300) for 1 hour at room temperature. Images were acquired using a Zeiss LSM900 Airyscan2 microscope and analysed using ImageJ plugins BIOPJACOB (interactions) or MitochondriaAnalyzer (morphology), as previously performed ^51^.

### Oxygen consumption rate (OCR) measurements

Cells were seeded in XFe24 Seahorse plates (Agilent, #100777-004) and allowed to attach for 36 hours. Cells were then grown overnight in complete DMEM or in glutamine-free DMEM. After treatments, cells were washed with PBS and incubated in DMEM-HG without glutamine (25 mM glucose, 1 mM pyruvate, 1% penicillin/streptomycin and 5 mM HEPES). Cells were allowed to equilibrate in an incubator without CO_2_ for 30 minutes. OCR were measured before and after acute addition of glutamine. Proton leak respiration was measured after the addition of 1 μM oligomycin and non-mitochondrial respiration after addition of 1 μM rotenone and 1 μM antimycin A.

### Mitochondrial H_2_O_2_ release

Cells (10^6^) were permeabilized as described above and incubated in 2 ml of experimental media (125 mM sucrose, 65 mM KCl, 10 mM HEPES, 2 mM KH_2_PO_4_, 2 mM MgCl_2_, 1 mM EGTA, and 0.1% BSA; pH 7.2) containing 5 U/mL horseradish peroxidase and 25 μM Amplex Red. The oxidation of Amplex Red generates a fluorescent compound known as resorufin, recorded on an F2500 Hitachi Fluorimeter at excitation and emission wavelengths of 563 and 587 nm, respectively ^10^.

### OMA1 activation protocol

Cells were grown in complete DMEM-HG for 24 hours. Cells were washed with PBS, and incubated in DMEM-HG without FBS in the presence of DMSO (Control), 20 µM Carbonyl Cyanide m-Chlorophenyl-hydrazone (CCCP) or 1 µM oligomycin plus 1 µM antimycin A.

## Statistical analysis of data

Statistical analyses were performed using Prism8 and Microsoft Excel. Data are expressed as mean ± SEM. Means were compared using Student’s *t*-test or ANOVA followed by Bonferroni post hoc test. Significance was accepted at p<0.05. Significance levels: p<0.05 (*); p<0.01 (**); p<0.001 (***); p<0.0001.

## Supporting information

supl mat

## Acknowledgments

This research was supported with funding from the Italian Ministry of Health (PRIN 20207P85MH to A.R.), European Union (Next-Generation EU CN00000041 to R.R), Fundação de Amparo à Pesquisa do Estado de São Paulo (FAPESP) grants 13/07937-8 and 20/06970-5, Conselho Nacional de Pesquisa e Desenvolvimento (CNPq), INCT and CEPID de Processos Redox em Biomedicina - Redoxoma, and Coordenação de Aperfeiçoamento de Pessoal de Nível Superior (CAPES) line 01. JDCS was supported by a FAPESP fellowships 19/05226-3 and 22/12251-7. We thank Sirley Mendes de Oliveira and Camille Caldeira da Silva for excellent technical support.

## Contributions

J.D.C.S., Investigation, Visualization, Writing – Original Draft and Conceptualization; D.D.A and G.O., Investigation; R.R., A.J.K. and A.R Resources, Writing – Review & Editing and Conceptualization.

---

**Supplementary Figure 1. Effects of starvation and refeeding on mitochondrial proteins in postprandial mouse liver.**

**(A)** – (**P**) Densitometric analyses for the levels of the MCUc components, p-S6, s-OPA1, CTF2, TMEM65, PDH and OXPHOS complexes of the Western blots shown in Fig. 1B and C, normalized as indicated. Data are presented as fold change relative to whole-liver lysates of animals fasted for 12 hours and represent the mean ± SEM. n≥3. For data analysis, one-way ANOVA was used with post hoc Bonferroni tests for each sample. * p≤0.05; ** p≤0.01; *** p≤0.001.

**Supplementary Figure 2. Effect of serum withdrawal and insulin on MCUc components.**

**(A)** – (**D**) Densitometric analyses for the levels of the MCUc components of the Western blot shown in Fig. 2B, normalized as indicated. Data are presented as fold change relative to -FBS and represent the mean ± SEM, n≥3. For data analysis, one-way ANOVA was used with post hoc Bonferroni tests for each sample. ** p≤0.01; *** p≤0.001.

**Supplementary Figure 3. Rapamycin modulates p-MICU1 levels.**

(**A**) Representative Western blots of HeLa cells cultured in control media (DMSO) or treated overnight with rapamycin probed with the indicated antibodies

(**B**) – (**E**) Densitometric analyses for the levels of the MCUc components of the Western blot shown in (A), normalized to β-actin expression. Data are presented as fold change relative to Control and represent the mean ± SEM. n≥3. For analysis, unpaired Student T-test was used. * p≤0.05; ** p≤0.01.

**Supplementary Figure 4. Glutamine deprivation reduces p-MICU1 levels, which can be prevented by proteasome inhibition.**

(**A**) Representative Western blots of HEK-293 cells cultured in control media (Control) or glutamine-starved conditions (-Gln) probed with the indicated antibodies

(**B**) – (**E**) Densitometric analyses for the levels of the MCUc components of the Western blot shown in (A), normalized to β-actin expression. Data are presented as fold change relative to Control and represent the mean ± SEM. n≥3. For analysis, unpaired Student T-test was used. * p≤0.05.

(**A**) Representative blots of HEK-293 cells cultured in glutamine-starved conditions in the presence (MG132) or absence (DMSO) of 10 μM proteasome inhibitor MG132 overnight probed with the indicated antibodies.

(**A**) – (**J**) Densitometric analyses for the levels of the MCUc components of the Western blot shown in (A), normalized to β-actin expression. Data are presented as fold change relative to DMSO and represent the mean ± SEM. n≥3. For analysis, unpaired Student T-test was used. * p≤0.05.

**Supplementary Figure 5. Glutamine starvation alters mitochondrial morphology, but not ER-mitochondria interactions.**

(**A**) HeLa cells mitochondria were labelled with anti-TOM20 and the ER with SEC61-β-GFP, in the presence (Control) or absence (-Gln) of glutamine. Co-localization, quantified by Mander’s coefficient 2 (fraction of ER surface overlapping mitochondria), was unchanged by glutamine deprivation. On the left, representative images. On the right, quantification of the experiment. n≥37. Data are presented as mean ± SEM.

(**B**) Mitochondrial morphology assessed by TOM20 immunofluorescence in HeLa cells in the presence (Control) or absence (-Gln) of glutamine. On the left, representative images. On the right, quantification of total branch length/mitochondrion, branches/mitochondrion, branch diameter, and Form Factor. Data are presented as mean ± SEM. n≥57. For analysis, unpaired Student T-test was used. * p≤0.05.

**Supplementary Figure 6. MICU1 turnover is independent of OMA1 and AFG3L2.**

HeLa cells were cultured in DMEM-HG without FBS under conditions promoting OMA1 activation: depolarization with 20 μM CCCP for 1 or 4 hours.

(**A**) Representative Western blots probed with the indicated antibodies. OPA1 was used as a readout of OMA1 activity: the upper bands disappear, and the lower bands increase upon depolarization.

(**B**) – (**D**) Densitometric analyses for the levels of the m-MICU1 (B), p-MICU1 (C) and i-MICU1

(**C**) of the Western blot shown in (A), normalized to β-actin expression. Data are presented as fold change relative to DMSO and represent the mean ± SEM. n≥3. For analysis, one-way ANOVA was used with post hoc Bonferroni tests for each sample.

(**D**) Cells were treated for 72 hours with non-targeting, AFG3L2, or OMA1 siRNAs. Representative Western blots probed with the indicated antibodies.

(**E**) and (**G**) Densitometric analyses for the levels of the MICU1 (F) and TMEM65 (G) of the Western blot shown in (E), normalized to β-actin expression. Data are presented as fold change relative to Control and represent the mean ± SEM. n≥3. For analysis, one-way ANOVA was used with post hoc Bonferroni tests for each sample. * p≤0.05; ** p≤0.01.

