## Supplementary material for "YME1L-dependent regulation of mitochondrial Ca^2+^ transport: a role for mitochondrial uptake protein 1 (MICU1) in nutrient sensing": supl mat

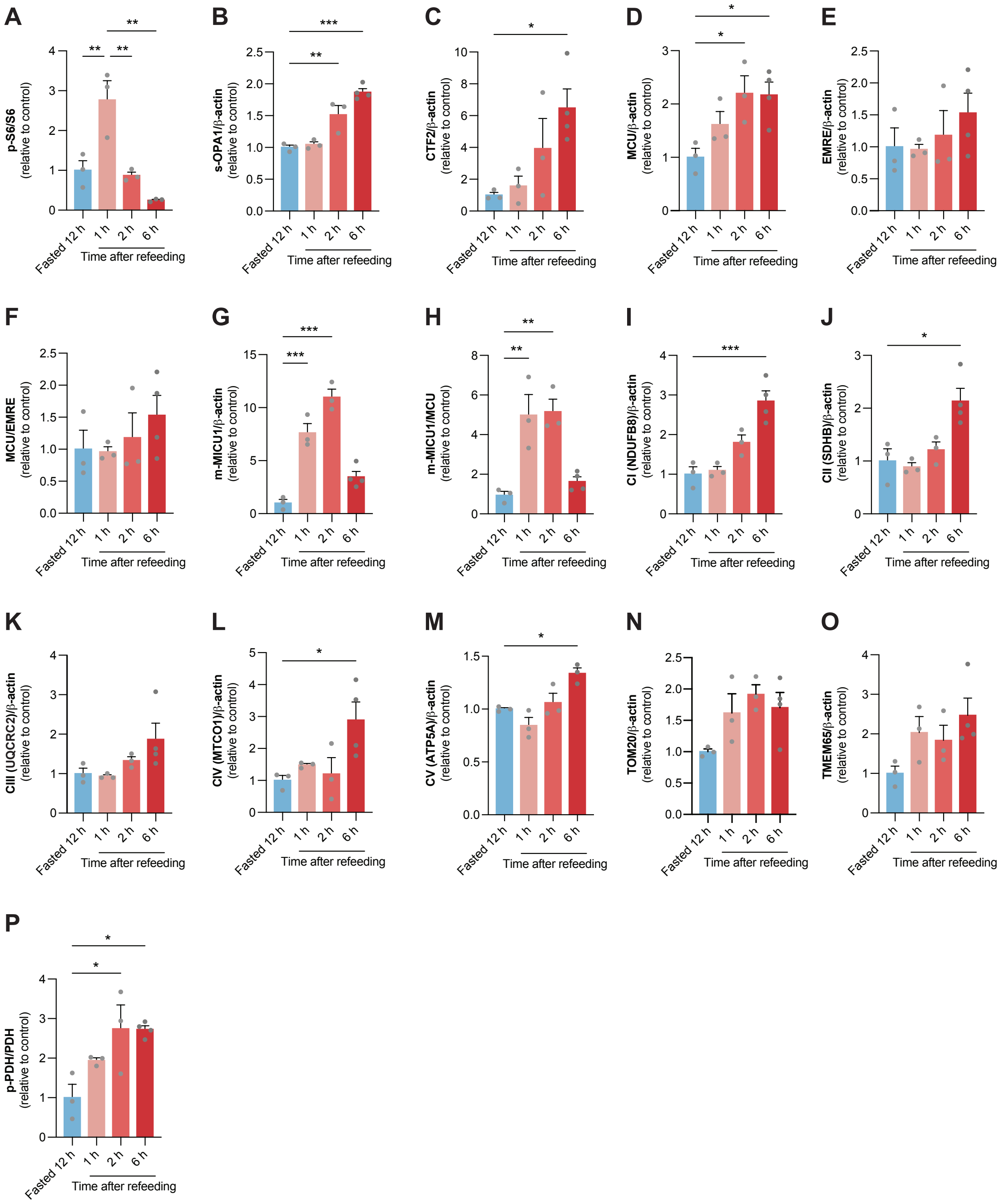


**Supplementary Figure 1. Effects of starvation and refeeding on mitochondrial proteins in postprandial mouse liver.**

(**A**) – (**P**) Densitometric analyses for the levels of the MCUc components, p-S6, s-OPA1, CTF2, TMEM65, PDH and OXPHOS complexes of the Western blots shown in Fig. 1B and C, normalized as indicated. Data are presented as fold change relative to whole-liver lysates of animals fasted for 12 hours and represent the mean ± SEM. n≥3. For data analysis, one-way ANOVA was used with post hoc Bonferroni tests for each sample. * p≤0.05; ** p≤0.01; *** p≤0.001.

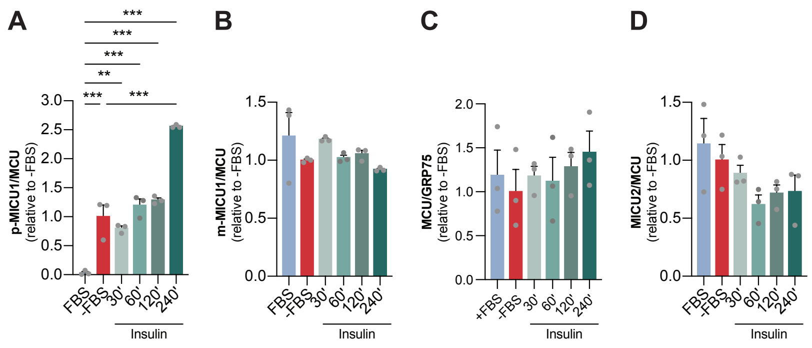


**Supplementary Figure 2. Effect of serum withdrawal and insulin on MCUc components**.

(**A**) – (**D**) Densitometric analyses for the levels of the MCUc components of the Western blot shown in Fig. 2B, normalized as indicated. Data are presented as fold change relative to -FBS and represent the mean ± SEM, n≥3. For data analysis, one-way ANOVA was used with post hoc Bonferroni tests for each sample. ** p≤0.01; *** p≤0.001.

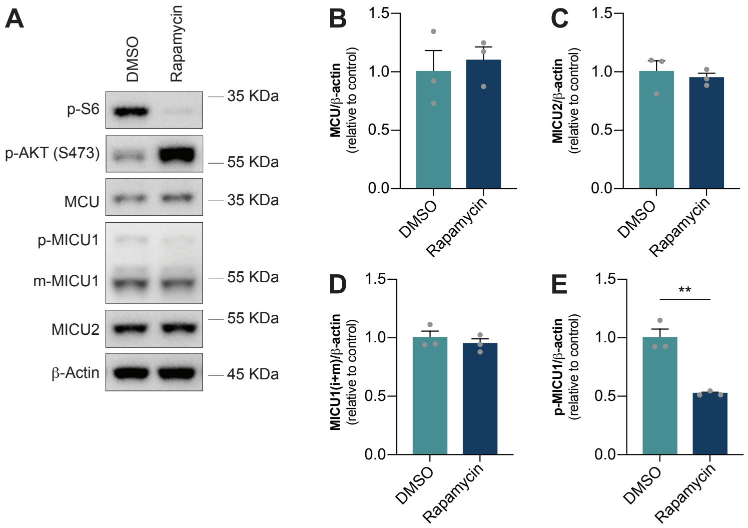


**Supplementary Figure 3. Rapamycin modulates p-MICU1 levels.**

(**A**) Representative Western blots of HeLa cells cultured in control media (DMSO) or treated overnight with rapamycin probed with the indicated antibodies
(**B**) – (**E**) Densitometric analyses for the levels of the MCUc components of the Western blot shown in (A), normalized to β-actin expression. Data are presented as fold change relative to Control and represent the mean ± SEM. n≥3. For analysis, unpaired Student T-test was used. * $p \leq 0.05$; ** $p \leq 0.01$.

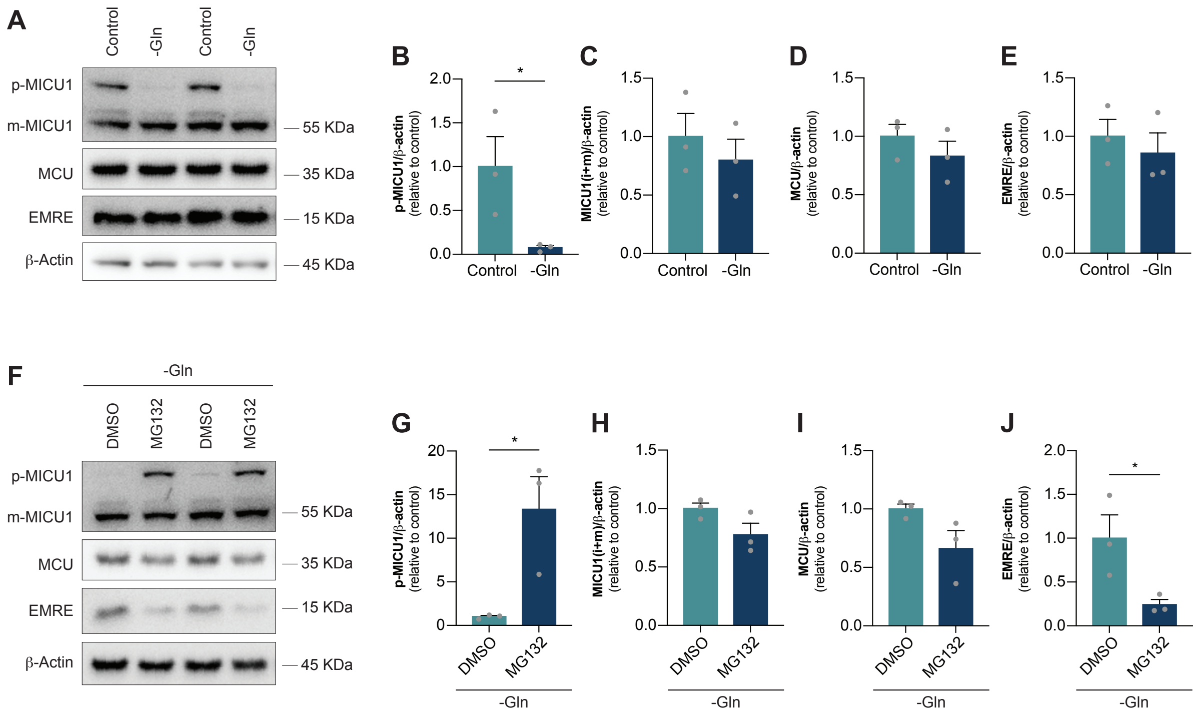


**Supplementary Figure 4. Glutamine deprivation reduces p-MICU1 levels, which can be prevented by proteasome inhibition.**

(**A**) Representative Western blots of HEK-293 cells cultured in control media (Control) or glutamine-starved conditions (-Gln) probed with the indicated antibodies

(**B**) – (**E**) Densitometric analyses for the levels of the MCUc components of the Western blot shown in (A), normalized to β-actin expression. Data are presented as fold change relative to Control and represent the mean ± SEM. n≥3. For analysis, unpaired Student T-test was used. * $p \leq 0.05$.

(**F**) Representative blots of HEK-293 cells cultured in glutamine-starved conditions in the presence (MG132) or absence (DMSO) of 10 μM proteasome inhibitor MG132 overnight probed with the indicated antibodies.

(**G**) – (**J**) Densitometric analyses for the levels of the MCUc components of the Western blot shown in (A), normalized to β-actin expression. Data are presented as fold change relative to DMSO and represent the mean ± SEM. n≥3. For analysis, unpaired Student T-test was used. * $p \leq 0.05$.

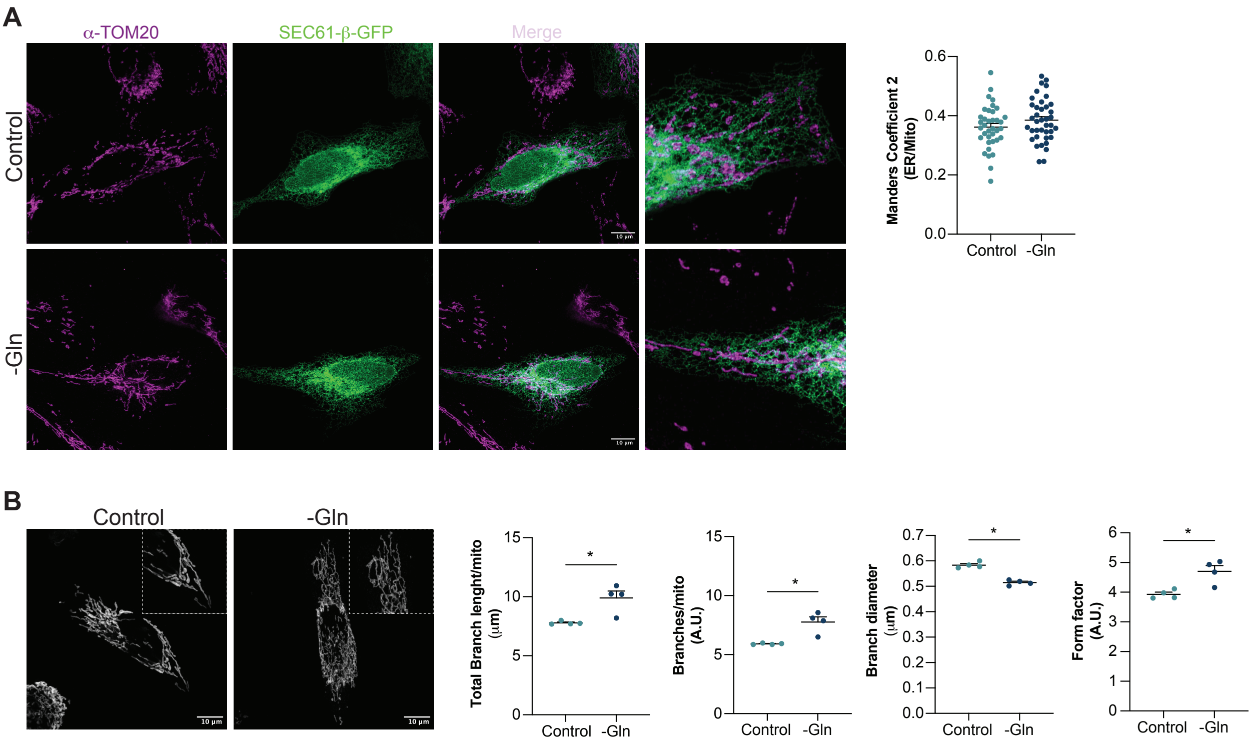


**Supplementary Figure 5. Glutamine starvation alters mitochondrial morphology, but not ER-mitochondria interactions.**

(**A**) HeLa cells mitochondria were labelled with anti-TOM20 and the ER with SEC61-β-GFP, in the presence (Control) or absence (-Gln) of glutamine. Co-localization, quantified by Mander´s coefficient 2 (fraction of ER surface overlapping mitochondria), was unchanged by glutamine deprivation. On the left, representative images. On the right, quantification of the experiment. n≥37. Data are presented as mean ± SEM.
(**B**) Mitochondrial morphology assessed by TOM20 immunofluorescence in HeLa cells in the presence (Control) or absence (-Gln) of glutamine. On the left, representative images. On the right, quantification of total branch length/mitochondrion, branches/mitochondrion, branch diameter, and Form Factor. Data are presented as mean ± SEM. n≥57. For analysis, unpaired Student T-test was used. * p≤0.05.

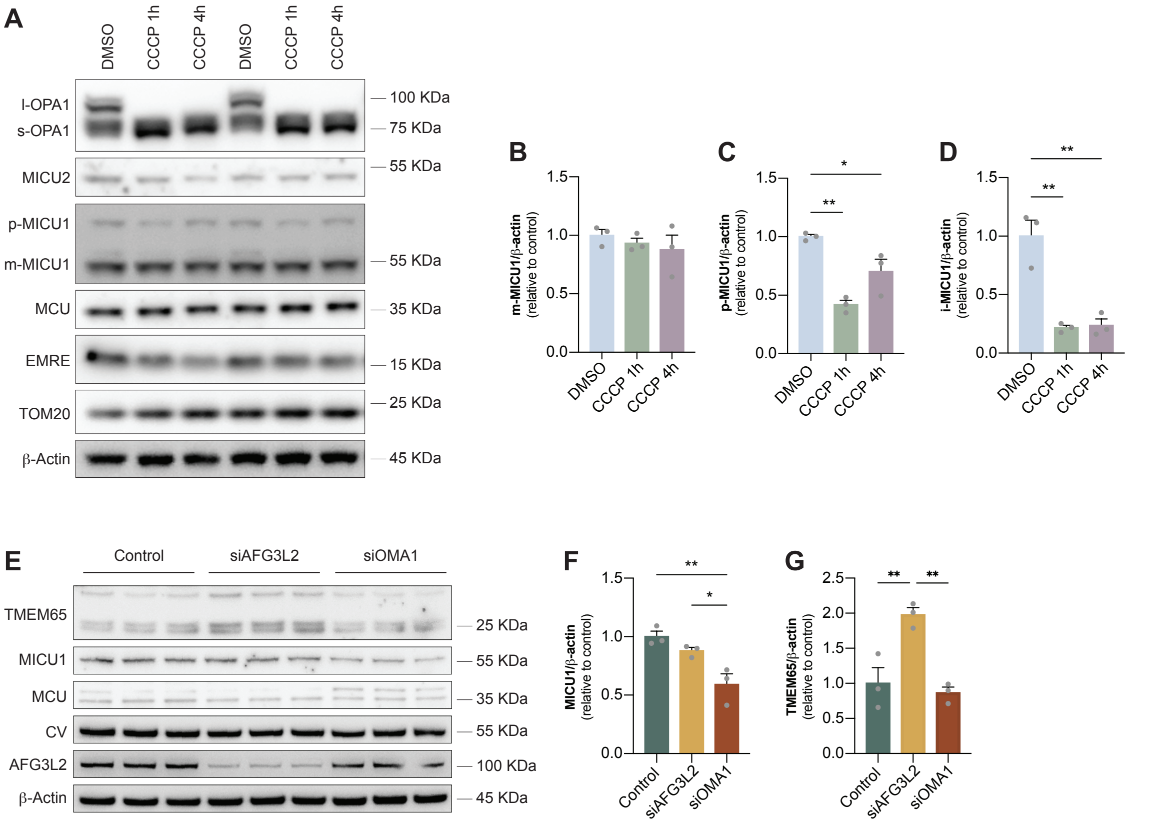


**Supplementary Figure 6. MICU1 turnover is independent of OMA1 and AFG3L2.**

HeLa cells were cultured in DMEM-HG without FBS under conditions promoting OMA1 activation: depolarization with 20 μM CCCP for 1 or 4 hours.

(**A**) Representative Western blots probed with the indicated antibodies. OPA1 was used as a readout of OMA1 activity: the upper bands disappear, and the lower bands increase upon depolarization.

(**B**) – (**D**) Densitometric analyses for the levels of the m-MICU1 (B), p-MICU1 (C) and i-MICU1 (D) of the Western blot shown in (A), normalized to β-actin expression. Data are presented as fold change relative to DMSO and represent the mean ± SEM. n≥3. For analysis, one-way ANOVA was used with post hoc Bonferroni tests for each sample.

(**E**) Cells were treated for 72 hours with non-targeting, AFG3L2, or OMA1 siRNAs. Representative Western blots probed with the indicated antibodies.

(**F**) and (**G**) Densitometric analyses for the levels of the MICU1 (F) and TMEM65 (G) of the Western blot shown in (E), normalized to β-actin expression. Data are presented as fold change relative to Control and represent the mean ± SEM. n≥3. For analysis, one-way ANOVA was used with post hoc Bonferroni tests for each sample. * $p \leq 0.05$; ** $p \leq 0.01$.
